# Spike mutation D614G alters SARS-CoV-2 fitness and neutralization susceptibility

**DOI:** 10.1101/2020.09.01.278689

**Authors:** Jessica A. Plante, Yang Liu, Jianying Liu, Hongjie Xia, Bryan A. Johnson, Kumari G. Lokugamage, Xianwen Zhang, Antonio E. Muruato, Jing Zou, Camila R. Fontes-Garfias, Divya Mirchandani, Dionna Scharton, John P. Bilello, Zhiqiang Ku, Zhiqiang An, Birte Kalveram, Alexander N. Freiberg, Vineet D. Menachery, Xuping Xie, Kenneth S. Plante, Scott C. Weaver, Pei-Yong Shi

## Abstract

A spike protein mutation D614G became dominant in SARS-CoV-2 during the COVID-19 pandemic. However, the mutational impact on viral spread and vaccine efficacy remains to be defined. Here we engineer the D614G mutation in the SARS-CoV-2 USA-WA1/2020 strain and characterize its effect on viral replication, pathogenesis, and antibody neutralization. The D614G mutation significantly enhances SARS-CoV-2 replication on human lung epithelial cells and primary human airway tissues, through an improved infectivity of virions with the spike receptor-binding domain in an “up” conformation for binding to ACE2 receptor. Hamsters infected with D614 or G614 variants developed similar levels of weight loss. However, the G614 virus produced higher infectious titers in the nasal washes and trachea, but not lungs, than the D614 virus. The hamster results confirm clinical evidence that the D614G mutation enhances viral loads in the upper respiratory tract of COVID-19 patients and may increases transmission. For antibody neutralization, sera from D614 virus-infected hamsters consistently exhibit higher neutralization titers against G614 virus than those against D614 virus, indicating that (i) the mutation may not reduce the ability of vaccines in clinical trials to protect against COVID-19 and (ii) therapeutic antibodies should be tested against the circulating G614 virus before clinical development.

**Importance:** Understanding the evolution of SARS-CoV-2 during the COVID-19 pandemic is essential for disease control and prevention. A spike protein mutation D614G emerged and became dominant soon after the pandemic started. By engineering the D614G mutation into an authentic wild-type SARS-CoV-2 strain, we demonstrate the importance of this mutation to (i) enhanced viral replication on human lung epithelial cells and primary human airway tissues, (ii) improved viral fitness in the upper airway of infected hamsters, and (iii) increased susceptibility to neutralization. Together with clinical findings, our work underscores the importance of this mutation in viral spread, vaccine efficacy, and antibody therapy.

## Introduction

Since the emergence of severe acute respiratory syndrome coronavirus 2 (SARS-CoV-2) in China in late 2019^1^, coronavirus disease 2019 (COVID-19) has caused >25 million confirmed infections and >850,000 fatalities worldwide. Hospitals and public health systems were overwhelmed first in Wuhan, followed by Italy, Spain, New York City, and other major cities, before cases peaked in these locations. Although most infections are mild, SARS-CoV-2 can cause severe, life-threatening pneumonia, particularly in older age groups and those with chronic pulmonary and cardiac conditions, diabetes, and other comorbidities. The exact mechanisms of severe disease remain unclear but typically involve a dysregulated, hyperinflammatory response following the initial stages of viral infection^2^. However, in addition to the host response, variation in viral strain phenotypes could also contribute to disease severity and spread efficiency.

Coronaviruses have evolved a genetic proofreading mechanism to maintain their long RNA genomes^3^. Despite the low sequence diversity of SARS-CoV-2^4^, mutations that mediate amino acid substitutions in the spike protein, which interacts with cellular receptors such as angiotensin-converting enzyme 2 (ACE2) to mediate entry into cells, can strongly influence host range, tissue tropism, and pathogenesis. During the SARS-CoV outbreak in 2002-2003, one such mutation was shown to mediate adaptation for infection of the intermediate civet host as well as for interhuman transmission^5^. For SARS-CoV-2, analyses of over 28,000 spike protein gene sequences in late May 2020 revealed a D614G amino acid substitution that was rare before March but increased in frequency as the pandemic spread^6^, reaching over 74% of all published sequences by June 2020^7^. The D614G substitution was accompanied by three other mutations: a C-to-T mutation in the 5’ untranslated genome region at position 241, a synonymous C-to-T mutation at position 3,037, and a nonsynonymous C-to-T mutation at position 14,408 in the RNA-dependent RNA polymerase gene^8^. This set of mutations not only increased globally, but during co-circulation within individual regions during outbreaks, suggesting a fitness advantage rather than simply founder effects or genetic drift. The association of spike protein amino acid substitutions with coronavirus transmissibility suggested that the D614G substitution was critical to this putative selective sweep. The correlation of this mutation with higher nasopharyngeal viral RNA loads in COVID-19 patients^6,9^ also supported a putative advantage of the mutant in transmission, which is key for viral fitness. However, direct measurements of fitness were needed to confirm this hypothesis.

Initial phenotypic characterizations of the D614G spike substitution were performed using pseudotyped viruses, whereby vesicular stomatitis virus (VSV) and lentiviral particles incorporating the SARS-CoV-2 spike protein alone were studied by replication kinetics. The production of significantly higher pseudotyped viral titers in multiple cell types by the G614 spike variant suggested that this substitution could be associated with enhanced entry into cells and replication in the airways of infected patients^6,7^. However, these results need to be confirmed in studies with authentic SARS-CoV-2 containing the spike 614 variant, and also using *in vivo* studies with a suitable animal model. Therefore, using an infectious cDNA clone for SARS-CoV-2^10^, we generated the D614G substitution in the January 2020 USA-WA1/2020 strain^11^ and performed experimental comparisons using *in vitro* cell culture, a primary human 3D airway tissue, and a hamster infection model^12^. We also developed a pair of D614 and G614 mNeonGreen SARS-CoV-2 viruses that could be used for rapid neutralization testing of serum specimens and monoclonal antibodies (mAbs). Using the reporter SARS-CoV-2 viruses, we analyzed the effect of D614G mutation on susceptibility to neutralization. Our study has important implications in understanding the evolution and transmission of SARS-CoV-2 as well as the development of COVID-19 vaccines and therapeutic antibodies.

## Results

### Enhancement of viral replication and infectivity by the spike D614G substitution in human lung epithelial cells

We first examined the effect of the spike D614G substitution on viral replication in cell culture. A site-directed mutagenesis was performed on an infectious cDNA clone of SARS-CoV-2 to prepare a pair of recombinant isogeneic viruses with spike D614 or G614 (Fig. 1a). Similar infectious amounts of D614 and G614 viruses were recovered from Vero E6 cells (monkey kidney epithelial cells), with viral titers of 1×10^8^ and 8×10^7^ plaque-forming units (PFU)/ml, respectively. The two viruses formed similar plaque morphologies (Fig. 1b). In Vero E6 cells, the G614 virus replicated to a higher infectious titer than D614 at 12 h post-infection (hpi), after which the two viruses replicated to comparable levels (Fig. 1c). A similar trend was observed for extracellular viral RNA production from the infected Vero E6 cells (Fig. 1d). To compare the infectivity between the two viruses, we calculated the genomic RNA/PFU ratios; no significant differences were found (Fig. 1e), indicating that the D614G mutation does not affect viral replication or virion infectivity on Vero E6 cells.

**Figure 1.**
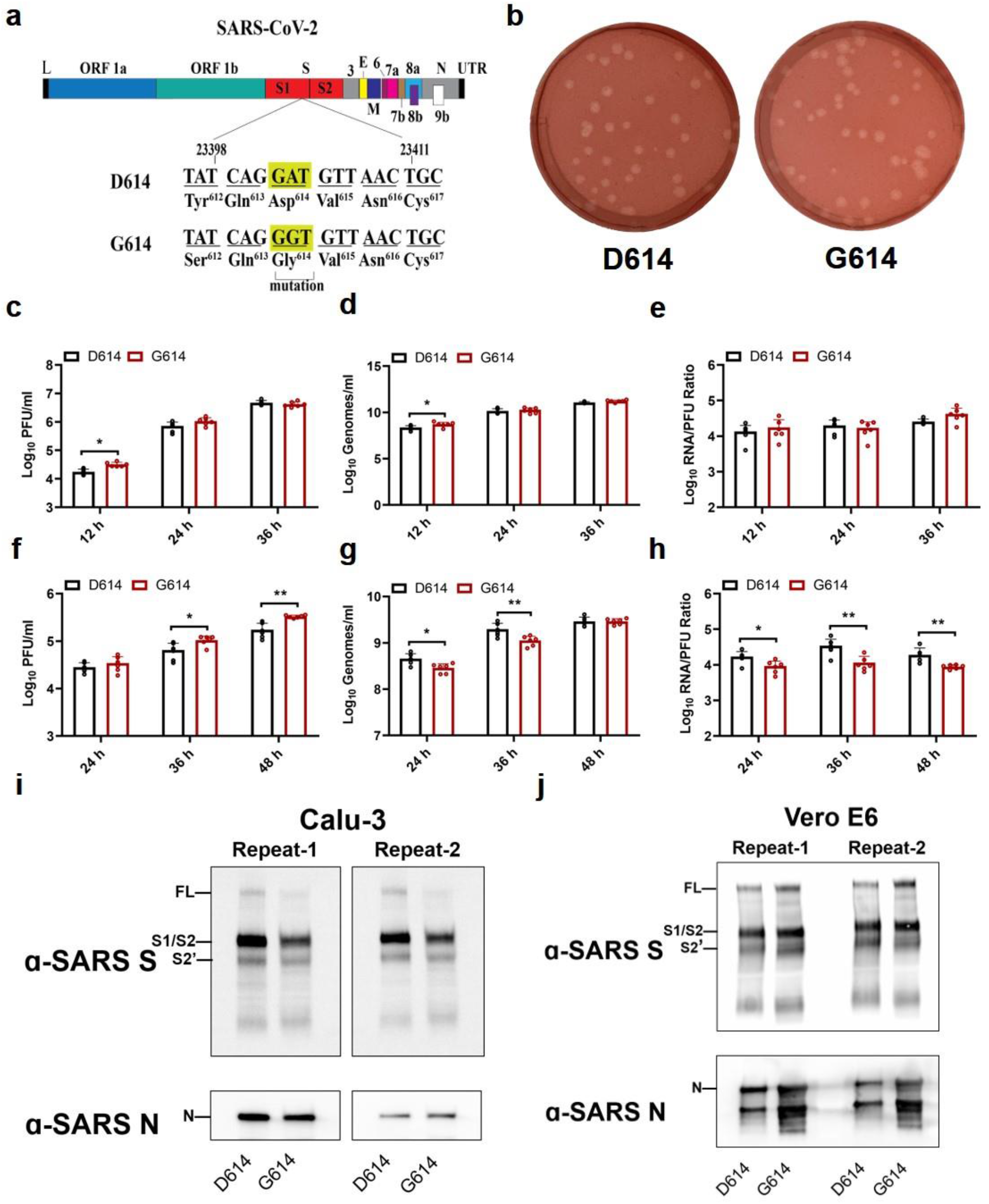
D614G substitution improves SARS-CoV-2 replication on Calu-3 cells through increased virion infectivity. **(a)** Construction of mutant G614 SARS-CoV-2. A single nucleotide A-to-G substitution was introduced to construct the spike D614G mutation in the infectious cDNA clone of SARS-CoV-2. The nucleotide positions of the viral genome are annotated. **(b)** Plaque morphologies of D614 and G614 viruses. The plaques were developed on day 2 post infection on Vero E6 cells. **(c-h)** Viral replication and genomic RNA/PFU ratios of D614 and G614 viruses produced from Vero E6 cells (c-e) and from Calu-3 cells (f-h). Both cells were infected with viruses at an MOI of 0.01. Infectious viral titers (c,f) and genomic RNA levels (d,g) in culture medium were determined by plaque assay and real-time RT-qPCR, respectively. The genomic RNA/PFU ratios (e,h) were calculated to indicate virion infectivity. The detection limitation of the plaque assay is 40 PFU/ml. The results were pooled from two independent biological replicates. Data are presented with mean ± standard deviations. *P* values were determined by two-tailed Mann–Whitney test. * *p*<0.05, ** *p*<0.01. **(i,j)** Spike protein cleavages of purified virions. Purified D614 and G614 virions were analyzed by Western blot using polyclonal antibodies against spike and anti-nucleocapsid antibodies. Full-length spike (FL), S1/S2 cleavage form, and S2’ protein are annotated. Results from two independent experiments are presented for virions produced from Calu-3 cells (i) and Vero E6 cells (j).

Next, we compared the replication kinetics of D614 and G614 viruses on the human lung epithelial Calu-3 cells. After infection at a multiplicity of infection (MOI) of 0.01 PFU/ml, the G614 virus produced modest 1.2-, 2.4-, and 1.9-fold more infectious virus than the D614 virus at 24, 36, and 48 hpi, respectively (Fig. 1f), indicating that D614G enhances viral replication. In contrast, the G614-infected cells produced less (at 24 and 36 hpi) or equivalent (at 48 hpi) extracellular viral RNA compared to D614-infected cells (Fig. 1g). The genomic RNA/PFU ratios of D614 virus were therefore 1.9-to 3.0-fold higher than those of G614 (Fig. 1h), indicating that the D614G mutation increases the infectivity of SARS-CoV-2 produced from the human lung cell line.

To explore the mechanism of increased infectivity of G614 virus produced from Calu-3 cells, we compared the spike protein processing from D614 and G614 viruses. Virions were purified from the culture medium of infected Calu-3 using ultracentrifugation and a sucrose cushion. The pelleted viruses were analyzed for spike protein processing by Western blot, with nucleocapsid included as a loading control. For both viruses, full-length spike was almost completely processed to the S1/S2 cleavage form and S2’, with comparable cleavage efficiencies of 93% for D614 and 95% for G614 (Fig. 1i). When virions produced from Vero E6 cells were analyzed, less full-length spike protein was processed to the S1/S2 form, with cleavage efficiencies of 73% for D614 and 67% for G614 (Fig. 1j). These results suggest that (i) more spike protein is cleaved to S1/S2 within virions produced from Calu-3 cells than those produced from Vero E6 cells and (ii) the D614G substitution does not significantly affect the spike cleavage ratio.

### Increased fitness in the hamster upper airway of SARS-CoV-2 with the D614G substitution

The *in vivo* relevance of the S-D614G mutation was evaluated in the golden Syrian hamster model (Extended Data Fig. 1a). After intranasally infecting four-to five-week-old hamsters with 2×10^4^ PFU of D614 or G614 virus, animals from both groups exhibited similar mean weight losses (Fig. 2a). No visible illness was observed in either infected cohort. On day 2 post-infection (pi), infectious viral titers from nasal washes, trachea, and various lobes of the lung (Extended Data Fig. 1b) were consistently higher in the G614-infected subjects compared to the D614-infected animals, although the differences did not reach statistical significance (Fig. 2b). The viral titer differences were greater in the upper airway samples than those in the lower airway tissues. On day 4 pi, the differences in infectious viral titers between the two viruses became more significant in the upper airway, with 1.3 log_10_ PFU/ml higher G614 virus than D614 in nasal wash (Fig. 2c). The trachea had 0.9 log_10_ PFU/g higher G614 virus than D614, but the statistical significance was lost upon correction for multiple comparisons. The viral loads in various lung lobes were nearly identical, with ≤0.1 log_10_ PFU/g differential between the two viruses. No infectious virus was detected in any airway tissues on day 7 pi (data not shown). Overall, the results demonstrate that the D614G mutation leads to a higher infectious virus production and shedding in the upper airway of infected hamsters.

**Figure 2.**
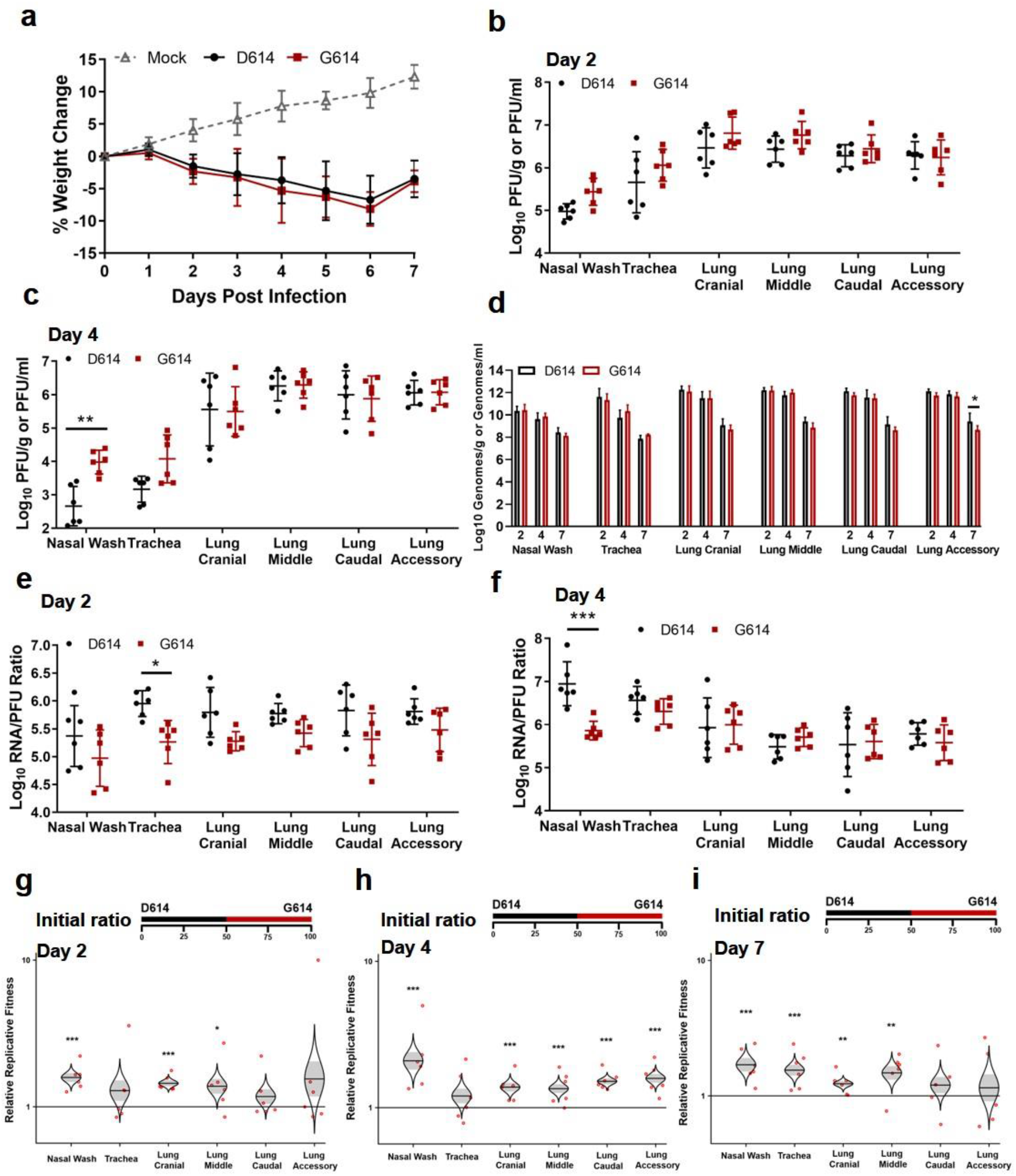
D614G substitution increases SARS-CoV-2 replication in the upper airway, but not the lungs, of hamsters. **(a)** Weight loss after infection with 2×10^4^ PFU of D614 or G614 SARS-CoV-2. Animals were also inoculated with PBS as a negative control. **(b,c)** Infectious titers in the nasal wash, trachea, and lung lobes on days 2 (b) and 4 (c) post infection. The infectious titers were quantified by plaque assay on Vero E6 cells. **(d)** Viral RNA levels on days 2, 4, and 7 post infection. The viral RNA levels were measured by RT-qPCR. **(e,f)** Viral RNA/PFU ratios on days 2 (e) and 4 (f) post infection. **(g-i)** Relative replicative fitness of D614 and G614 viruses on days 2 (g), 4 (h), and 7 (i) post infection. Data reflect 6 mice per infected cohort for each timepoint. For weight loss (a), symbols represent the mean. For infectious titers and viral RNA/PFU ratios (b,c,e,f), symbols represent individual animals and midlines represent the mean. For viral RNA levels (d), bar height represents the mean. For all graphs, error bars represent standard deviations. Weight loss (a) was analyzed by two-factor ANOVA with virus strain and timepoint as fixed factors, and a T ukey’s post-hoc test to compare all cohort pairs at a given timepoint. All other datasets (b-f) were analyzed by measures two-factor ANOVA with virus strain and tissue as fixed factors, and a Sidak’s post-hoc test to compare D614 versus G614 within a given tissue. For the competition assay (g-i), 100 μl mixtures of equal D614 and G614 viruses (10^4^ PFU per virus) were inoculated intranasally into 4-to 5-week-old Syrian hamsters. The initial ratio of D614 and G614 viruses is 1:1. Organs of infected hamsters were collected on days 2 (g), 4 (h), and 7 (i) post infection and measured for the relative fitness of G614 virus over D614 virus using Sanger sequencing. The distribution of the model-adjusted means is illustrated by catseye plots with shaded +/- standard error overlaid by scatterplots of subject measures; scatterplots have been randomly jittered horizontally for clarity, and are shown on the log_10_ scale such that comparisons are a null value of 1. Reported p-values are based on the results of the respective post-hoc tests. * *p*<0.05, ** *p*<0.01, *** *p*<0.001.

We compared the infectivity of the D614 and G614 viruses produced in hamsters by determining their viral RNA levels and viral RNA/PFU ratios. In contrast to the higher infectious titers for G614 than D614 virus, the two viruses produced nearly identical levels of viral RNA across all organs and timepoints (Fig 2d). The RNA/PFU ratios of G614 virus were 0.3 log_10_ to 0.7 log_10_ lower than those of D614 virus across airway tissues (Fig. 2e). On day 4 pi, a 1.1 log_10_ lower RNA/PFU ratio was detected for G614 than D614 in nasal wash, while the differences in the trachea and lungs were 0.1 log_10_ to 0.3 log_10_ (Fig. 2f). On day 7 pi., despite no detectable infectious virus (detection limit 40 PFU/ml), more than 10^8^ viral RNA copies/ml were detected in the nasal washes (Fig. 2d), demonstrating high levels of viral RNA persistence after the clearance of infectious virus; this result recapitulates findings in COVID-19 patients, who frequently tested positive with RT-PCR for up to several weeks but have low or undetectable infectious virus. One caveat of the above RNA/PFU calculation was that the total RNA could include viral RNAs from both virions and lysed cells during sample processing. Nevertheless, the results suggest that mutation D614G may enhance the infectivity of SARS-CoV-2 in the respiratory tract, particularly in the upper airway of infected animals.

The above results prompted us to directly compare the finesses of D614 and G614 viruses through a competition experiment. This approach has major advantages over performing individual strain infections with numerous host replicates; each competition is internally controlled, eliminating host-to-host variation that can reduce the power of experiments, and the virus strain ratios can be assayed with more precision than individual virus titers. Thus, competition assays have been used for many studies of microbial fitness, including viruses^13–16^. To perform the competition between D614 and G614 variants, we intranasally infected hamsters with equal amounts of the two viruses (10^4^ PFU per virus). Since the infecting viruses were prepared from Vero E6 cells with comparable viral RNA/PFU ratios (Fig. 1e), the animals also received equivalent levels of D614 and G614 viral RNA. On days 2, 4, and 7 pi, nasal wash and respiratory organs were harvested and quantified for relative amounts of D614 and G614 RNAs by RT-PCR and Sanger sequencing. The ratios of G614/D614 RNA were then calculated from the electropherograms to indicate the relative fitness for viral replication. A G614/D614 ratio of >1.0 or <1.0 indicates a replication advantage for G614 or D614, respectively. As shown in Fig. 2h-i, all respiratory tissues showed G614/D614 ratios of 1.2 to 2.6 on days 2, 4, and 7 pi, indicating that G614 virus has a consistent advantage over D614 virus when infecting the respiratory tract of hamsters.

### Dramatic enhancement of viral replication by spike mutation D614G in a primary human airway tissue model

To further define the function of D614G mutation in human respiratory tract, we characterized the replication of D614 and G614 viruses in a primary human airway tissue model (Fig. 3a). This airway model contains human tracheal/bronchial epithelial cells in multilayers which resemble the epithelial tissue of the respiratory tract. The primary tissue is cultured at an air-liquid interface to recapitulate the barrier, microciliary response, and infection of human airway tissues *in vivo*^17,18^. After infecting the airway tissue at an MOI of 5, both D614 and G614 viruses produced increasing infectious titers from day 1 to 5, up to 7.8×10^5^ PFU/ml (Fig. 3b), demonstrating that the airway tissue supports SARS-CoV-2 replication. The infectious viral titers of G614 were significantly higher (2.1-to 8.6-fold) than those of D614 (Fig. 3b). In contrast, no differences in viral RNA yields were observed between the two viruses (Fig. 3c). The genomic RNA/PFU ratios of D614 virus were 1.4-to 5.3-fold higher than those of G614 virus (Fig. 3d). Sequencing of the D614 and G614 viruses collected on day 5 pi did not show any other mutations. Collectively, the results demonstrate that substitution D614G enhances viral replication through increased virion infectivity when SARS-CoV-2 replicates on primary human upper airway tissues.

**Figure 3.**
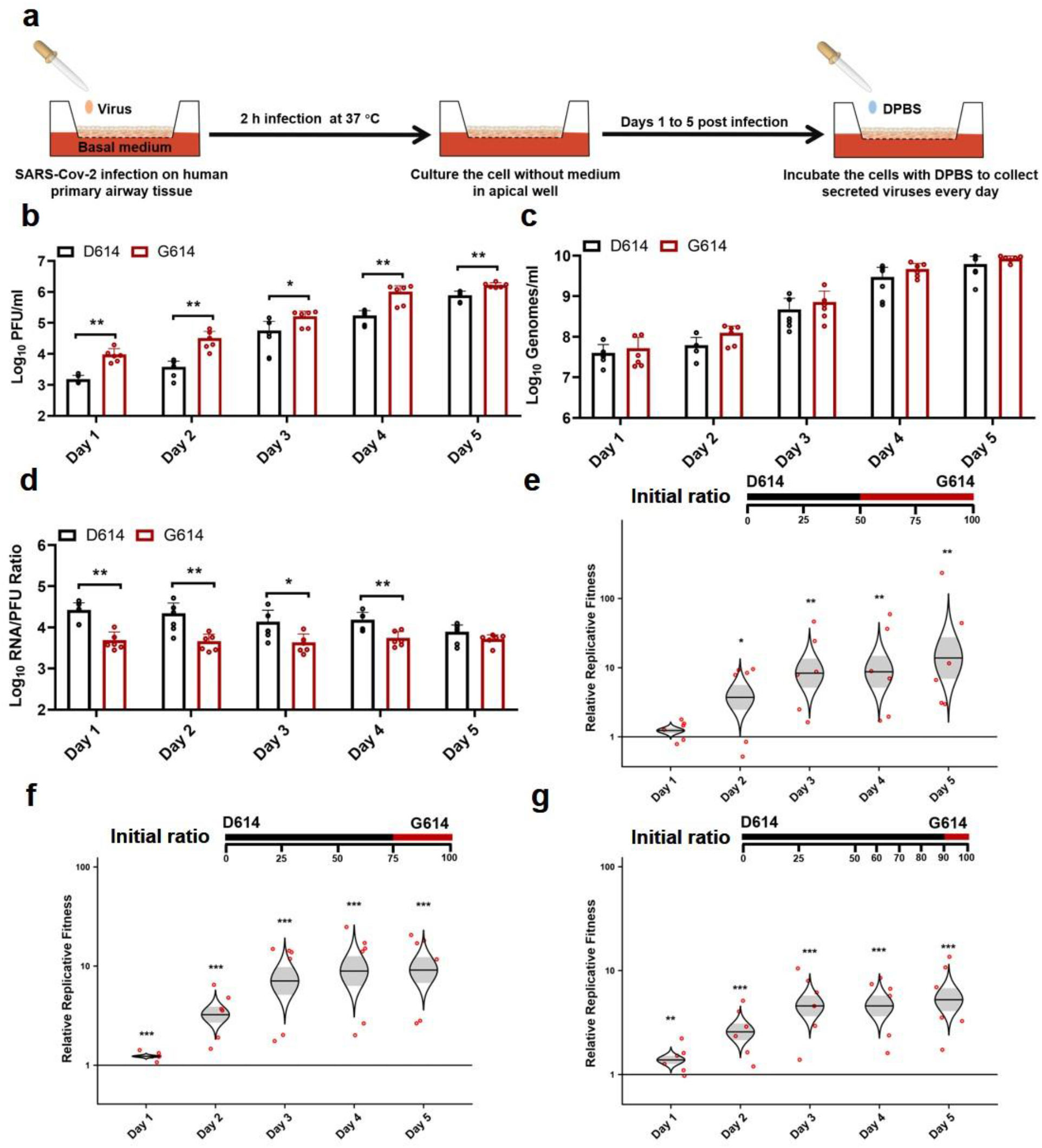
D614G substitution significantly enhances SARS-CoV-2 replication in primary human airway tissues. **(a)** Experimental scheme. D614 and G614 viruses were inoculated onto the primary human airway tissues. After incubation at 37°C for 2 h, the culture was washed with DPBS for three times to remove the un-attached virus. The culture was maintained at 37°C, 5% CO_2_ for 5 days. On each day, 300 μl DPBS was added onto the culture. After incubation at 37°C for 30 min, the DPBS containing the eluted viruses was subjected to plaque assay, real-time RT-qPCR, and competition analysis by Sanger sequencing. **(b-d)** Viral replication and genomic RNA/PFU ratios. Human airway tissues were infected with D614 or G614 virus at an MOI of 5. The amounts of infectious virus (b) and genomic RNA (c) were quantified by plaque assay and real-time RT-qPCR, respectively. The genomic RNA/PFU ratio (d) was calculated to indicate virion infectivity. The results were pooled from two independent biological replicates. Data are presented as means ± standard deviations. *P* values were determined by two-tailed Mann–Whitney test. **(e,f)** Competition assay. A mixture of D614 and G614 viruses with different initial ratios were inoculated onto the human airway tissues at a total MOI of 5. The initial D614/G614 virus ratio was 1:1 (e), 3:1(f), or 9:1(g). The G614/D614 ratios after competition were measure by Sanger sequencing and analyzed using R statistical software. The distribution of the model-adjusted means is illustrated by catseye plots with shaded +/- standard error (SD) overlaid by scatterplots of subject measures; scatterplots have been randomly jittered horizontally for clarity, and are shown on the log_10_ scale such that comparisons are a null value of 1. \**p* < 0.05, ** *p*< 0.01, *** *p*< 0.001.

Next, we performed competition experiments to directly compare the replication fitness of D614 and G614 viruses in the human airway culture. After infecting with a 1:1 infectious ratio of D614 and G614 viruses (produced from Vero E6 cells), the G614/D614 ratios increased to 1.2, 3.7, 8.2, 8.8, and 13.9 on days 1, 2, 3, 4, and 5 pi, respectively (Fig. 3e). In addition, after infecting the airway culture with 3:1 ratio of D614 and G614 viruses, the G614 variant was able to rapidly overcome its initial deficit to reach a slight advantage with a G614/D614 ratio of 1.2 by day 1 pi, with that advantage increasing to 9.1-fold by day 5 pi (Fig. 3f). Furthermore, when infecting the airway tissue with 9:1 ratio of D614 and G614 viruses, the G614/D614 ratios increased from 1.4 to 5.2 from days 1 to 5 pi (Fig. 3g). A similar competition result was obtained when the experiment was repeated using a different donor-derived human airway culture (Extended Data Fig. 2). These competition results confirm that the G614 virus can rapidly outcompete the D614 virus when infecting human airway tissues, even initially as a minor variant in a mixed population.

### Effect of spike mutation D614G on neutralization susceptibility

All of the COVID-19 vaccines currently in clinical trials are based on the original D614 spike sequence^19,20^. An important question is whether substitution D614G could reduce vaccine efficacy, assuming G614 virus continues to circulate. To address this question, we measured the neutralization titers of a panel of sera collected from hamsters that were previously infected with D614 SARS-CoV-2 (Extended Data Fig. 3). Each serum was analyzed by a pair of mNeonGreen reporter SARS-CoV-2 viruses with the D614 or G614 spike (Extended Data Fig. 4)^21^. The mNeonGreen gene was engineered at the open-reading-frame 7 of the SARS-CoV-2 genome^10^. As shown in Figs. 4a-c, all sera exhibited 1.4-to 2.3-fold higher neutralization titers (mean 1.7-fold) against G614 virus than those against D614 (Extended Data Fig. 5), suggesting that mutation D614G may confer higher susceptibility to serum neutralization.

**Figure 4.**
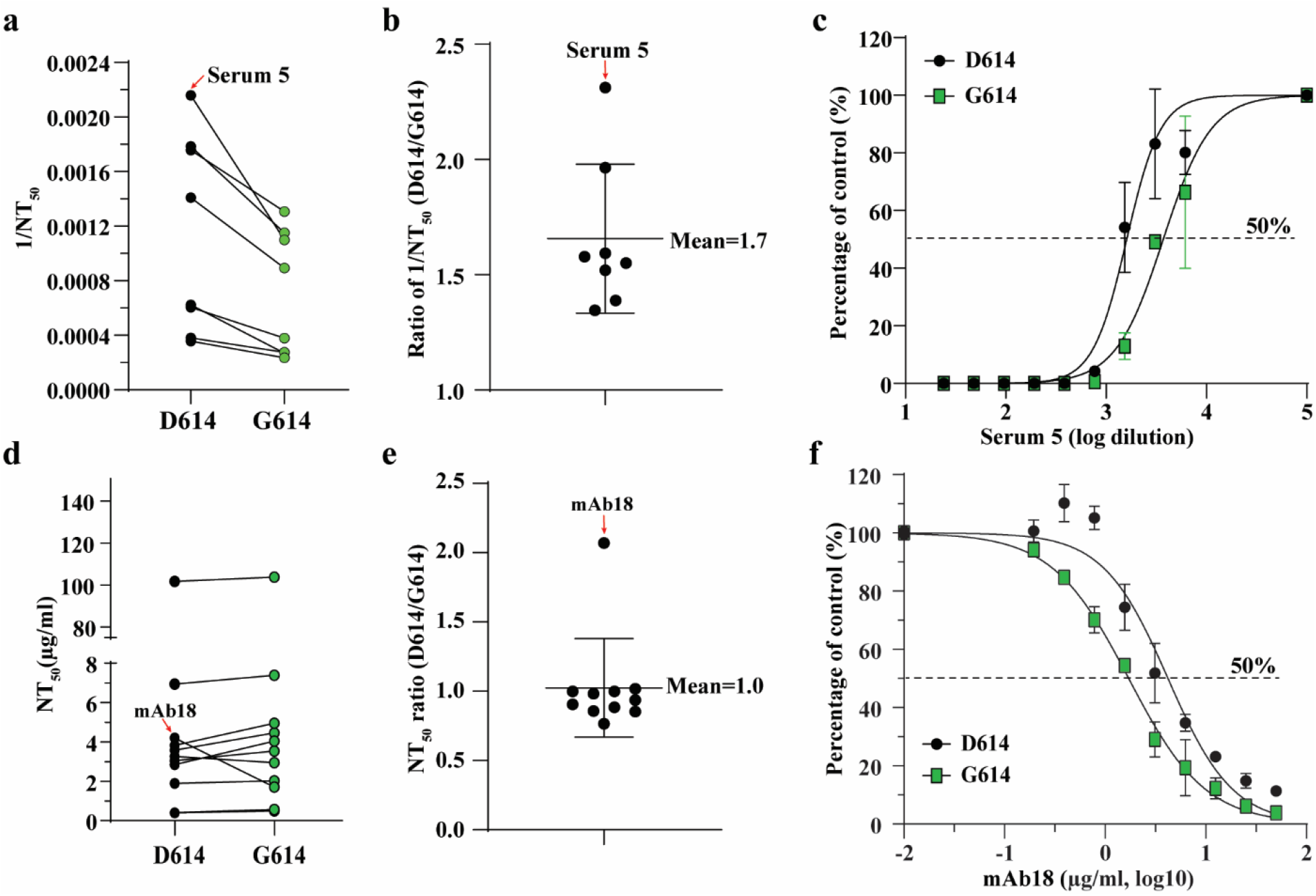
D614G substitution affects the neutralization susceptibility of SARS-CoV-2 to neutralizing sera and mAbs. **(a)** Neutralizing activities of hamster sera against D614 and G614 mNeonGreen reporter SARS-CoV-2. Eight sera from D614 virus-infected hamsters were tested for neutralizing titers against D614 and G614 reporter SARS-CoV-2. The 1/NT_50_ values for individual sera are plotted. **(b)** Ratio of 1/NT_50_ between D614 and G614 viruses. The mean of the ratios [ratio = (D614 1/NT_50_)/(G614 1/NT_50_)] from 8 hamster serum samples are shown. Error bar indicates the standard deviations. **(c)** Representative neutralizing curve of hamster serum 5. Dot line indicates 50% inhibition of viral infection. The means and standard deviations from two replicates are shown. **(d)** Neutralizing activities of eleven human mAbs against D614 and G614 mNeonGreen SARS-CoV-2. The data represents one of the two independent experiments. **(c)** Ratio of NT_50_ between D614 and G614 viruses. The averages of the NT_50_ ratios from two independent experiments performed in duplicates are shown. The mean and standard deviation from eleven mAbs are indicated. **(f)** Representative neutralizing curve of mAb18. Dotted line indicated 50% viral inhibition. The means and standard deviations from two replicates are shown.

To further examine the role of D614G mutation in antibody recognition and neutralization, we evaluated a panel of eleven human receptor-binding domain (RBD) mAbs against the D614 and G614 mNeonGreen SARS-CoV-2 viruses. The details of these RBD mAbs are reported elsewhere (An et al., submitted for publication). One mAb (mAb18) showed a 2.1-fold higher potency against G614 than D614 virus, whereas the other ten mAbs exhibited similar neutralization activities against both viruses (Figs. 4d-f and Extended Data Figs. 6 and 7). The results suggest that mutation D614G may modulate spike protein conformation to affect mAb neutralization in an epitope-specific manner.

## Discussion

We demonstrated that the spike substitution D614G enhanced SARS-CoV-2 replication in the upper respiratory tract through increased virion infectivity. Compared with the original D614 virus, the emergent, now dominant G614 virus replicated to a higher level in the human lung Calu-3 cells and primary human upper airway tissues. The replication differences were more dramatically observed in the human airway culture, with up to a 13.9-fold advantage when the two viruses were compared in a head-to-head competition test. The increased replication fitness corelated with an enhanced specific infectivity of the G614 virion. Since previous studies with pseudotyped virus showed that the cleavage efficiency of the spike protein into S1/S2 modulates SARS-CoV-2 infection^22,23^, we compared the S1/S2 ratios between the D614 and G614 virions. Although virions produced from Calu-3 cells had more complete S1/S2 cleavage than those produced form Vero E6 cells, no substantial differences in spike cleavage were detectable between the D614 and G614 virions produced from either cell type, suggesting that the enhanced virion infectivity is not likely due to the D614G-mediated spike cleavage difference. Our results from authentic SARS-CoV-2 are in contrast with previous studies reporting that the D614G mutation changes the cleavage and shedding of spike protein when expressed alone or in the context of pseudotyped virions^24,25^. Mechanistically, two recent studies showed that the D614G mutation abolishes a hydrogen-bond interaction with T859 from a neighboring protomer of the spike trimer^6^, which allosterically promotes the RDB domain to an “up” conformation for receptor ACE2 binding and fusion^7^, leading to an enhanced virion infectivity.

The higher viral loads of G614 in the upper airway of COVID-19 patients^26^ and infected hamsters support the role of D614G mutation in viral transmissibility. The robust replication of SARS-CoV-2 in the human upper airway may be partially conferred by a higher ACE2 receptor expression in the nasal cavity compared to that in the lower respiratory tract^27,28^. Patients infected with G614 virus developed higher levels of viral RNA in the nasopharyngeal swabs than those infected with D614 virus, but disease severity is not associated with the D614G mutation^6,7,9^. Our hamster infection model recapitulated these clinical findings: The G614 virus developed higher infectious titers than the D614 virus in nasal washes and tracheas, but not lungs; no differences in weight loss or signs of disease were observed between the G614- and D614-infected animals. If the lower viral RNA/PFU ratio of the G614 virus observed in our hamster and human airway models could be extrapolated to COVID-19 patients, the modest differences in cycle threshold (Ct) values of RT-qPCR observed in patients’ nasopharyngeal swabs would translate to ?10-fold infectious G614 virus, underscoring the potential for enhanced transmission and spread. This potential is further bolstered by the observation that a COVID-19 patient with two distinct populations of SARS-CoV-2 in the throat swabs and sputum samples only transmitted the throat strain to individuals downstream in the transmission chain^26,29^. Similar nasal-driven transmission was recently reported for human influenza A virus in a ferret model^30^.

Our results showed that G614 virus is consistently more susceptible to neutralization by sera collected from D614 virus-infected hamsters. The increased susceptibility of G614 virus to serum neutralization generated by D614 seems counterintuitive, but could be explained by the D614G-mediated increase in the “up” conformation of the RDB for binding to ACE2 receptor^6,7^. Since current COVID-19 vaccines in clinical trials are based on the original D614 sequence, our neutralization result mitigates the concern that the D614G mutation might compromise the efficacy of vaccines against the circulating G614 virus. Future studies are needed to eliminate this concern by testing human sera collected from the D614 spike vaccinatees. Besides antisera, we also showed that, depending on the epitope locations on RBD, the neutralizing potency of certain mAbs may be affected by the D614G mutation. The results underscore the importance to test therapeutic mAbs against G614 and other newly emerged mutant viruses during preclinical development.

In summary, we have used authentic SARS-CoV-2 to demonstrate that spike substitution D614G enhances viral replication in the upper respiratory tract and increases neutralization susceptibility. These findings have important implications in understanding the evolution and spread of the ongoing COVID-19 pandemic, vaccine efficacy, and therapeutic antibody development.

## Methods

### Ethics statement

Hamster studies were performed in accordance with the guidance for the Care and Use of Laboratory Animals of the University of Texas Medical Branch (UTMB). The protocol was approved by the Institutional Animal Care and Use Committee (IACUC) at UTMB. All the hamster operations were performed under anesthesia by isoflurane to minimize animal suffering.

### Animals and Cells

The Syrian hamsters (HsdHan:AURA strain) were purchased from Envigo (Indianapolis, IN). African green monkey kidney epithelial Vero E6 cells were grown in Dulbecco’s modified Eagle’s medium (DMEM) with 5% fetal bovine serum (FBS; HyClone Laboratories, South Logan, UT) and 1% antibiotic/ streptomycin (Gibco). Human lung adenocarcinoma epithelial Calu-3 2B4 cells were maintained in a high-glucose DMEM supplemented with 10% FBS and 1% penicillin/streptomycin at 37°C with 5% CO_2_. The EpiAirway system is a primary human airway 3D tissue model purchased from MatTek Life Science (Ashland, MA). This EpiAirway system was maintained with the provided culture medium at 37°C with 5% CO_2_ following the manufacturer’s instruction. All other culture medium and supplements were purchased from ThermoFisher Scientific (Waltham, MA). All cell lines were verified and tested negative for mycoplasma.

### Generation of SARS-CoV-2 spike D614G mutant viruses

One single-nucleotide substitution was introduced into a subclone puc57-CoV-2-F5-7 containing the spike gene of the SARS-CoV-2 wild type (WT) infectious clone^10^ to convert the 614^th^ amino acid from aspartic acid (D) to glycine (G) by overlap fusion PCR. The full-length infectious cDNA clone of SARS-CoV-2 D614G was assembled by *in vitro* ligation of seven contiguous cDNA fragments following the protocol previously described^10^. For construction of D614G mNeonGreen SARS-CoV-2, seven SARS-CoV-2 genome fragments (F1 to F5, F6 containing D614G mutation, and F7-mNG containing the mNeonGreen reporter gene) were prepared and *in vitro* ligated as described previously^10^. *In vitro* transcription was then preformed to synthesize full-length genomic RNA. For recovering the mutant viruses, the RNA transcripts were electroporated into Vero E6 cells. The viruses from electroporated cells were harvested at 40 h post electroporation and served as seed stocks for subsequent experiments. The D614G mutation from the recovered viruses was confirmed by sequence analysis. Viral titers were determined by plaque assay on Vero E6 cells. All virus preparation and experiments were performed in a biosafety level 3 (BSL-3) facilities.

### RNA extraction, RT-PCR, and Sanger sequencing

Cell culture supernatants or clarified tissue homogenates were mixed with a five-fold excess of TRIzol™ LS Reagent (Thermo Fisher Scientific, Waltham, MA). Viral RNAs were extracted according to the manufacturer’s instructions. The extracted RNAs were dissolved in 20 μl nuclease-free water. Two microliters of RNA samples were used for reverse transcription by using the SuperScript™ IV First-Strand Synthesis System (ThermoFisher Scientific) with random hexamer primers. Nine DNA fragments flanking the entire viral genome were amplified by PCR. The resulting DNAs were cleaned up by the QIAquick PCR Purification Kit, and the genome sequences were determined by Sanger sequencing at GENEWIZ (South Plainfield, NJ).

The quantify viral RNA samples, quantitative real-time RT-PCR assays were performed using the iTaq SYBR Green One-Step Kit (Bio-Rad) on the LightCycler 480 system (Roche, Indianapolis, IN) following the manufacturers’ protocols. Primers are listed in Extended Data Table 1. The absolute quantification of viral RNA was determined by a standard curve method using an RNA standard (*in* vitro transcribed 3,839bp containing genomic nucleotide positions 26,044 to 29,883 of SARS-CoV-2 genome).

To quantify D614:G614 ratios for competition assays, a 663-bp RT-PCR product was amplified from extracted RNA using a SuperScript™ III One-Step RT-PCR kit (Invitrogen, Carlsbad, CA, USA). A 20-μl reaction was assembled in PCR 8-tube strips through the addition of 10 μl 2× reaction mix, 0.4 μl SuperScript III RT/Platinum Taq Mix, 0.8 μl Forward Primer (10 μM) (Extended Data Table 1), 0.8μl reverse primer (10 μM) (Extended Data Table 1), 4 μl RNA, and 6 μl Rnase-free water. Reverse transcription and amplification was completed using the following protocol: (i) 55°C, 30 min; 94°C, 2 min; (ii) 94°C, 15 s; 60°C, 30 s; 68°C, 1 min; 40 cycles; (iii) 68°C, 5 min; (iv) indefinite hold at 4°C. The presence and size of the desired amplicon was verified with 2 μl of PCR product on an agarose gel. The remaining 18 μl were purified by a QIAquick PCR Purification kit (Qiagen,Germantown, MD) according to the manufacturer’s protocol.

Sequences of the purified RT-PCR products were generated using a BigDye Terminator v3.1 cycle sequencing kit (Applied Biosystems, Austin, TX, USA). The sequencing reactions were purified using a 96-well plate format (EdgeBio, San Jose, CA, USA) and analyzed on a 3500 Genetic Analyzer (Applied Biosystems, Foster City, CA).The peak electropherogram height representing each mutation site and the proportion of each competitor was analyzed using the QSVanalyser program^31^.

### Plaque assay

Approximately 1.2×10^6^ Vero E6 cells were seeded to each well of 6-well plates and cultured at 37°C, 5% CO_2_ for 16 h. Virus was serially diluted in either DMEM with 2% FBS (for viral stocks and *in vitro*-generated samples) or DPBS (for hamster tissues) and 200 μl was transferred to the monolayers. The viruses were incubated with the cells at 37°C with 5% CO_2_ for 1 h. After the incubation, overlay medium was added to the infected cells per well. The overlay medium contained either DMEM with 2% FBS and 1% sea-plaque agarose (Lonza, Walkersville, MD) in the case of *in vitro* samples or Opti-MEM with 2% FBS, 1% penicillin/streptomycin, and 0.8% agarose in the case of *in vivo* samples. After a 2-day incubation, plates were stained with neutral red (Sigma-Aldrich, St. Louis, MO) and plaques were counted on a light box.

### Viral infection on cells

Approximately 3×10^5^ Vero E6 or Calu-3 cells were seeded onto each well of 12-well plates and cultured at 37°C, 5% CO_2_ for 16 h. Either SARS-CoV-2 D614 or G614 virus was inoculated into the cells at an MOI of 0.01. The virus was incubated with the cells at 37°C for 2 h. After the infection, the cells were washed by DPBS for 3 times to remove the unattached virus. One milliliter of culture medium was added into each well for the maintenance of the cells. At each time point, 100 μl of culture supernatants were harvested for the real-time qPCR detection and plaque assay. Meanwhile, 100 μl fresh medium was added into each well to replenish the culture volume. The cells were infected in triplicates for each virus. All samples were stored in −80°C freezer until plaque or RT-PCR analysis.

### Virion purification and spike protein cleavage analysis

Vero E6 or Calu-3 2B4 cells were infected with D614 or G614 viruses at an MOI of 0.01. At 24 (for Vero) or 48 (Calu-3) hpi, the culture media were collected and clarified by low speed spin. Virions in the media were pelleted by ultracentrifugation through a 20% sucrose cushion at 26,000 rpm for 3 h at 4°C by in a Beckman SW28 rotor. The purified virions were analyzed by Western blot using polyclonal antibodies against spike protein and nucleocapsid as described previously^32^.

### Viral infection in a primary human airway tissue model

The EpiAirway system is a primary human airway 3D mucociliary tissue model consisting of normal, human-derived tracheal/bronchial epithelial (HAE) cells. For viral replication kinetics study, either D614 or G614 virus was inoculated onto the culture at an MOI of 5 in DPBS. After 2 h infection at 37°C with 5% CO_2_, the inoculum was removed, and the culture was washed three times with DPBS. The infected epithelial cells were maintained without any medium in the apical well, and medium was provided to the culture through the basal well. The infected cells were incubated at 37°C, 5% CO_2_. From day 1 to day 5, 300 μl DPBS were added onto the apical side of the airway culture and incubated at 37°C for 30 min to elute the released viruses. All virus samples in DPBS were stored at −80°C.

### Hamster infection

Four-to five-week-old male golden Syrian hamsters, strain HsdHan:AURA (Envigo, Indianapolis, IN), were inoculated intranasally with 2×10^4^ PFU SARS-CoV-2 in a 100-μl volume. Eighteen animals received WT D614 virus, 18 received mutant G614 virus, and 18 received a mixture containing 10^4^ PFU of D614 virus and 10^4^ PFU of G614 virus. The infected animals were weighed and monitored for signs of illness daily. On days 2, 4, and 7 pi, cohorts of 6 infected animals and 4 (days 2 and 4) or 6 (day 7) mock-infected animals were anesthetized with isoflurane and nasal washes were collected in 400 μl sterile DPBS. Animals were humanely euthanized immediately following the nasal wash. The trachea and the four lobes of the right lung were harvested in maintenance media (DMEM supplemented with 2% FBS and 1% penicillin/streptomycin) and stored at −80°C. Samples were subsequently thawed, tissues were homogenized for 1 min at 26 sec-1, and debris was pelleted by centrifugation for 5 min at 16,100×g. Infectious titers were determined by plaque assay. Genomic RNAs were quantified by quantitative RT-PCR (Extended Data Table 1). Ratios of D614/G614 RNA were determined via RT-PCR with quantification of Sanger peak heights.

### Competition assay

For the competition on primary human airway 3D tissue model, the D614 and G614 mutant viruses were mixed and inoculated onto the cells at a final MOI of 5. The initial ratio of D614 and G614 viruses is 1:1, 3:1, or 9:1 based on PFU titers determined on Vero E6 cells. The DPBS with viruses was harvested every day from day 1 to 5 following the protocol described above. For the competition in hamsters, 100 μl mixtures of D614 and G614 viruses (total 2×10^4^ PFU per hamster) were inoculated intranasally into 4-5 weeks old Syrian hamsters. On days 2, 4, and 7 pi, 6 infected hamsters were sampled for competition detection. An aliquot of the inoculum for both hamster and human airway infections was back titered for estimating the initial ratio of viruses. All samples were stored in −80°C freezer prior to analysis.

### Neutralization assay

Neutralization assays were preformed using D614 and G614 mNeonGreen SARS-CoV-2 as previously described^21^. Briefly, Vero (CCL-81) cells were plated in black μCLEAR flat-bottom 96-well plate (Greiner Bio-one™). On the following day, sera or monoclonal antibodies were serially diluted from 1/20 starting dilution and nine 2-fold dilutions to the final dilution of 1/5,120 and incubated with D614 or G614 mNeonGreen SARS-CoV-2 at 37°C for 1 h. The virus-serum mixture was transferred to the Vero cell plate with the final MOI of 2.0. After 20 h, Hoechst 33342 Solution (400-fold diluted in Hank’s Balanced Salt Solution; Gibco) was added to stain cell nucleus, sealed with Breath-Easy sealing membrane (Diversified Biotech), incubated at 37°C for 20 min, and quantified for mNeonGreen fluorescence using Cytation™ 7 (BioTek). The raw images (2×2 montage) were acquired using 4× objective, processed, and stitched using the default setting. The total cells (indicated by nucleus staining) and mNeonGreen-positive cells were quantified for each well. Infection rates were determined by dividing the mNeonGreen-positive cell number to total cell number. Relative infection rates were obtained by normalizing the infection rates of serum-treated groups to those of non-serum-treated controls. The curves of the relative infection rates versus the serum dilutions (log_10_ values) were plotted using Prism 8 (GraphPad). A nonlinear regression method was used to determine the dilution fold that neutralized 50% of mNeonGreen fluorescence (NT_50_). Each serum was tested in duplicates.

### Statistics

Male hamsters were randomly allocated into different groups. The investigators were not blinded to allocation during the experiments or to the outcome assessment. No statistical methods were used to predetermine sample size. Descriptive statistics have been provided in the figure legends. For *in vitro* replication kinetics, Kruskal–Wallis analysis of variance was conducted to detect any significant variation among replicates. If no significant variation was detected, the results were pooled for further comparison. Differences between continuous variables were assessed with a non-parametric Mann–Whitney test. Hamster weights were analyzed by two factor ANOVA, with the percent weight change as the dependent variable and the strain and time as fixed factors. Tukey’s post-hoc test was used to compare all cohort pairs on days 1-7 pi. Log_10_-tranformed titers were analyzed by two-factor repeated measures ANOVA with the organ and strain as fixed factors. Sidak’s post-hoc test was used to compare strains within each organ. Genomic RNA/PFU ratios were calculated from non-transformed values, and the resulting ratios were log_10_-transformed prior to two factor repeated measures ANOVA with the organ and strain as fixed factors and Sidak’s post-hoc test to compare strains within a given organ. When a sample was below the limit of detection, it was treated as half of the limit of detection value for statistical and graphing purposes. Analysis was performed in Prism version 7.03 (GraphPad, San Diego, CA).

For virus competition experiments, relative replicative fitness values for G614 strain over D614 strain were analyzed according to w=(f0/i0), where i0 is the initial D614/G614 ratio and f0 is the final D614/G614 ratio after competition. Sanger sequencing (initial timepoint T0) counts for each virus strain being compared were based upon average counts over three replicate samples of inocula per experiment, and post-infection (timepoint T1) counts were taken from samples of individual subjects. For the primary human airway samples, multiple experiments were performed, so that f0/i0 was clustered by experiment. To model f0/i0, the ratio T0/T1 was found separately for each subject in each strain group, log (base-10) transformed to an improved approximation of normality, and modeled by analysis of variance with relation to group, adjusting by experiment when appropriate to control for clustering within experiment. Specifically, the model was of the form Log_10__CountT1overCountT0 ~ Experiment + Group. Fitness ratios between the two groups [the model’s estimate of w=(f0/i0)] were assessed per the coefficient of the model’s Group term, which was transformed to the original scale as 10^coefficient. This modeling approach compensates for any correlation due to clustering within experiment similarly to that of corresponding mixed effect models, and is effective since the number of experiments was small. Statistical analyses were performed using R statistical software (R Core Team, 2019, version 3.6.1). In all statistical tests, two-sided alpha=.05. Catseye plots^33^, which illustrate the normal distribution of the model-adjusted means, were produced using the “catseyes” package^34^.

## Acknowledgments

This research was supported by grants from NIA and NIAID of the NIH [AI153602 and AG049042 to V.D.M.; R24AI120942 (WRCEVA) to S.C.W] and by STARs Award provided by the University of Texas System to V.D.M. P.-Y.S. was supported by NIH grants AI142759, AI134907, AI145617, and UL1TR001439, and awards from the Sealy & Smith Foundation, Kleberg Foundation, the John S. Dunn Foundation, the Amon G. Carter Foundation, the Gilson Longenbaugh Foundation, and the Summerfield Robert Foundation. A.M. is supported by a Clinical and Translational Science Award NRSA (TL1) Training Core (TL1TR001440) from NIH. J.L. and C.R.F.-G. were supported by the McLaughlin Fellowship at the University of Texas Medical Branch.

## Author contributions

Conceptualization, Y.L., V.D.M., X.X., K.S.P., S.C.W., P.-Y.S.; Methodology, J.A.P, Y.L., J.L., H.X., B.A.J., K.G.L., X.Z., A.E.M., J.Z., C.R.F.G., A.N.F., V.D.M., X.X., K.S.P., S.C.W., P.- Y.S.; Investigation, J.A.P, Y.L., J.L., H.X., B.A.J., K.G.L., X.Z., A.E.M., J.Z., C.R.F.G., D.M., D.S., J.P.B., A.N.F., V.D.M., X.X., K.S.P., S.C.W., P.-Y.S.; Resources, Z.K., Z.A.; Data Curation, J.A.P, Y.L., J.L., H.X., B.A.J., K.G.L., X.Z., A.E.M., J.Z., A.N.F., V.D.M., X.X., K.S.P. S.C.W., P.-Y.S.; Writing-Original Draft, J.A.P, Y.L., J.L., X.X., K.S.P., S.C.W., P.-Y.S; Writing-Review & Editing, J.A.P, Y.L., J.L., H.X., B.A.J., K.G.L., X.Z., A.E.M., J.Z., A.N.F., V.D.M., X.X., K.S.P., S.C.W., P.- Y.S.; Supervision, A.N.F., V.D.M., X.X., K.S.P., S.C.W., P.-Y.S.; Funding Acquisition, A.N.F., V.D.M., S.C.W., P.-Y.S..

---

**Extended Data Figure 1.**
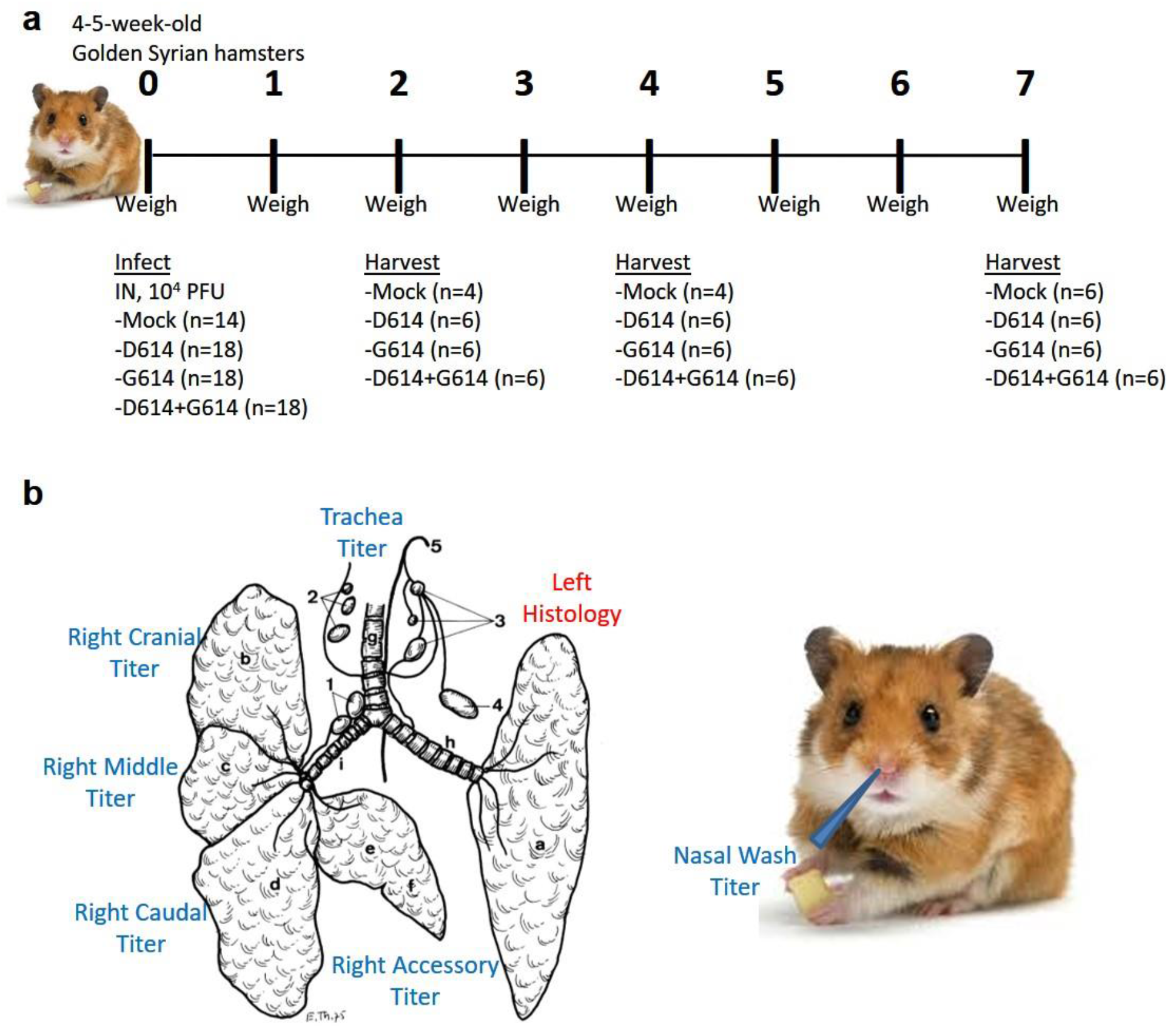
Experimental design of hamster infection and sample harvest. **(a)** Graphical overview of experiment to assess the impact of G614 mutation on replication in the respiratory system of hamsters. **(b)** Schematic samples harvested on days 2, 4, and 7 post-infection. Illustration of hamster lung adapted from Reznik, G. *et al. Clinical anatomy of the European hamster. Cricetus cricetus, L*., For sale by the Supt of Docs, U.S. Govt. Print. Off., 1978.

**Extended Data Figure 2.**
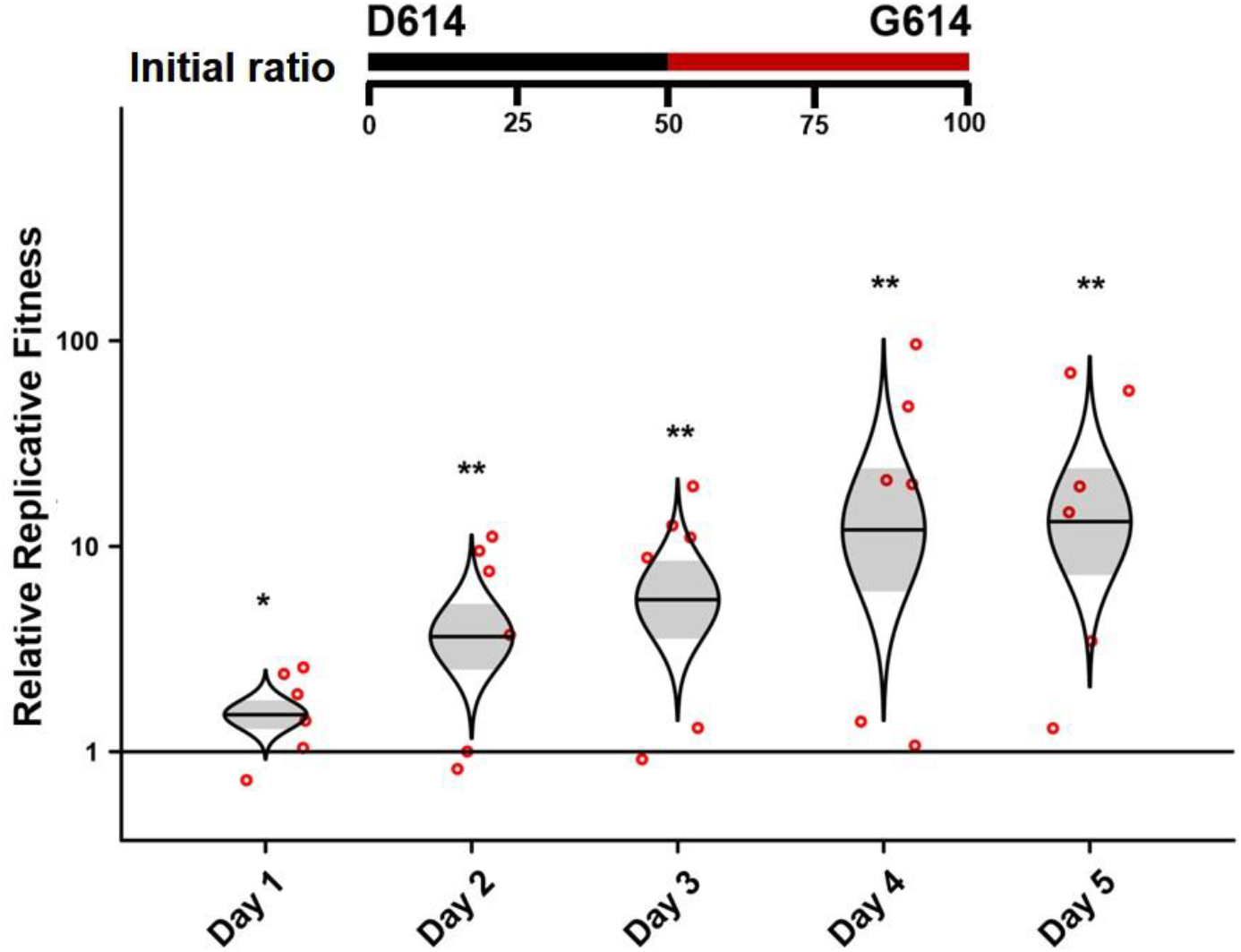
D614G substitution significantly enhances SARS-CoV-2 replication in primary human airway tissues from a different donor. D614 and G614 viruses were equally mixed and inoculated onto the airway tissue at a total MOI of 5. This airway tissue was produced from a different donor that was used in Figure 3. The tissues were washed by DPBS to collect the secreted viruses every day from days 1 to 5. The total RNAs were isolated and amplified by RT-PCR. The ratio of D614 and G614 viruses after competition were measure by Sanger sequencing and analyzed using R statistical software. The distribution of the model-adjusted means is illustrated by catseye plots with shaded +/- standard error (SD) overlaid by scatterplots of subject measures; scatterplots have been randomly jittered horizontally for clarity, and are shown on the log (base-10) scale such that comparisons are a null value of 1. \**p* < 0.05, ** *p*< 0.01.

**Extended Data Figure 3.**
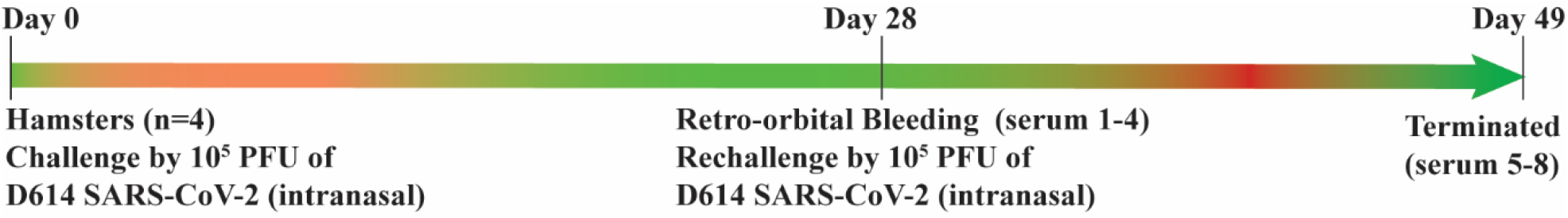
Scheme for preparing the D614 SARS-CoV-2-infected hamster sera for neutralization assay. Eight sera were collected: Four sera (number 1-4) collected on day 28 post infection and another four sera (number 5-8) collected on day 49 after the second viral infection.

**Extended Data Figure 4.**
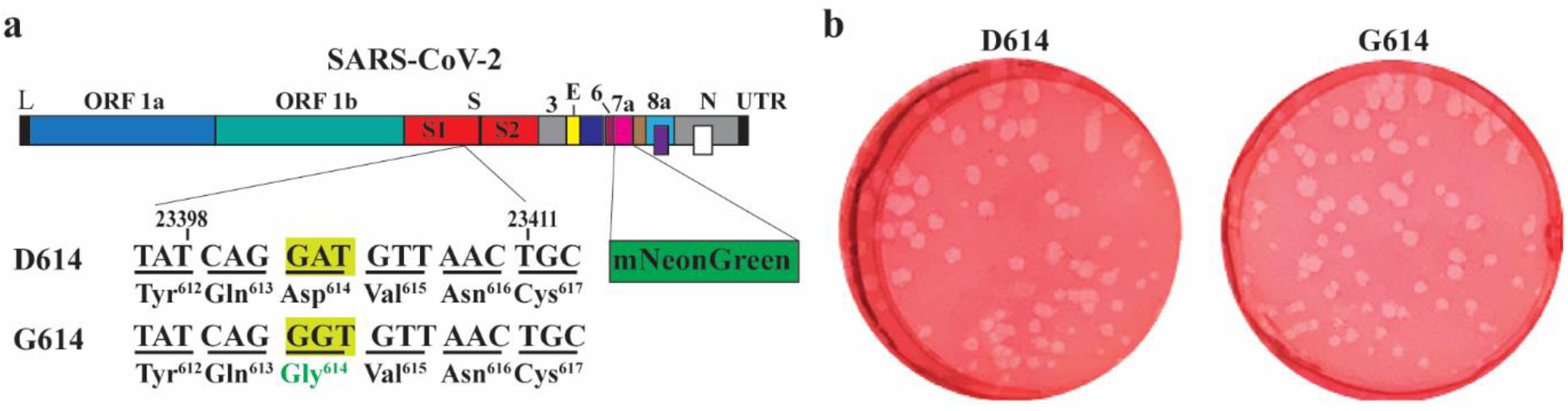
Construction of G614 mNeonGreen SARS-CoV-2. **(a)** Diagram of the construction. The D614G mutation was introduced into a mNeonGreen reporter SARS-CoV-2 using the method as described previously^10^. **(b)** Plaque morphologies of D614 and G614 mNeonGreen SARS-CoV-2.

**Extended Data Figure 5.**
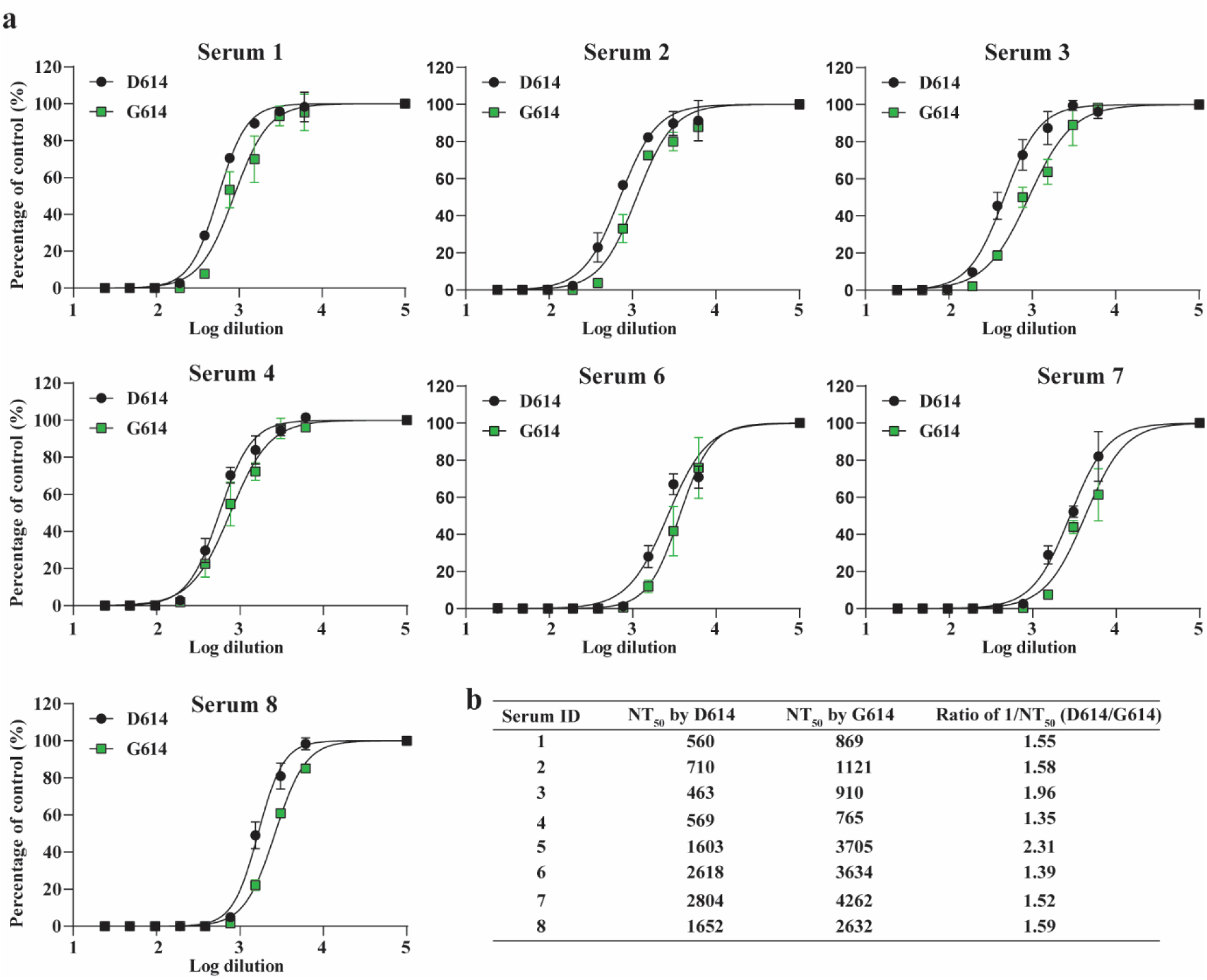
Neutralization activities of hamster sera against D614 and G614 mNeonGreen SARS-CoV-2. **(a)** Neutralizing curves of eight hamster sera against D614 and G614 mNeonGreen SARS-CoV-2. The neutralizing curve for serum 5 is shown in Fig. 4c. Experiments were performed in replicates. The mean and standard deviations are shown. **(b)** Calculated NT_50_ values and ratios of 1/NT_50_ for all eight hamster sera. The mean ratios were determined by (D614 1/NT_50_)/(G614 1/NT_50_).

**Extended Data Figure 6.**
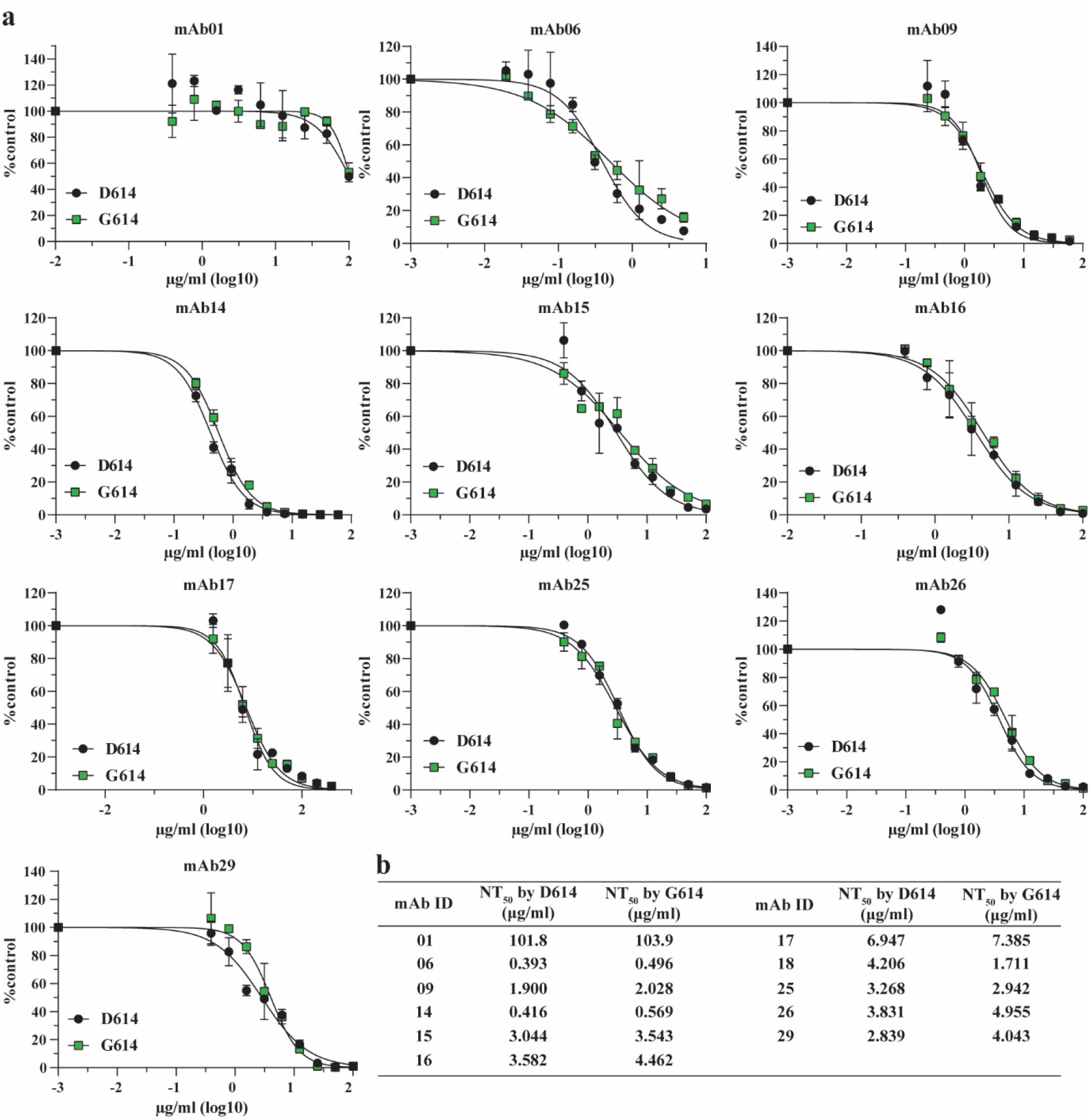
Neutralization activities of human mAbs against D614 and G614 mNeonGreen SARS-CoV-2 in Experiment I. **(a)** Neutralizing curves of eleven mAbs against D614 and G614 reporter SARS-CoV-2. The neutralizing curve for mAb18 is shown in Fig. 4f. Experiments were performed in replicates. The mean and standard deviations are shown. **(b)** Calculated NT_50_ values for all eleven mAbs.

**Extended Data Figure 7.**
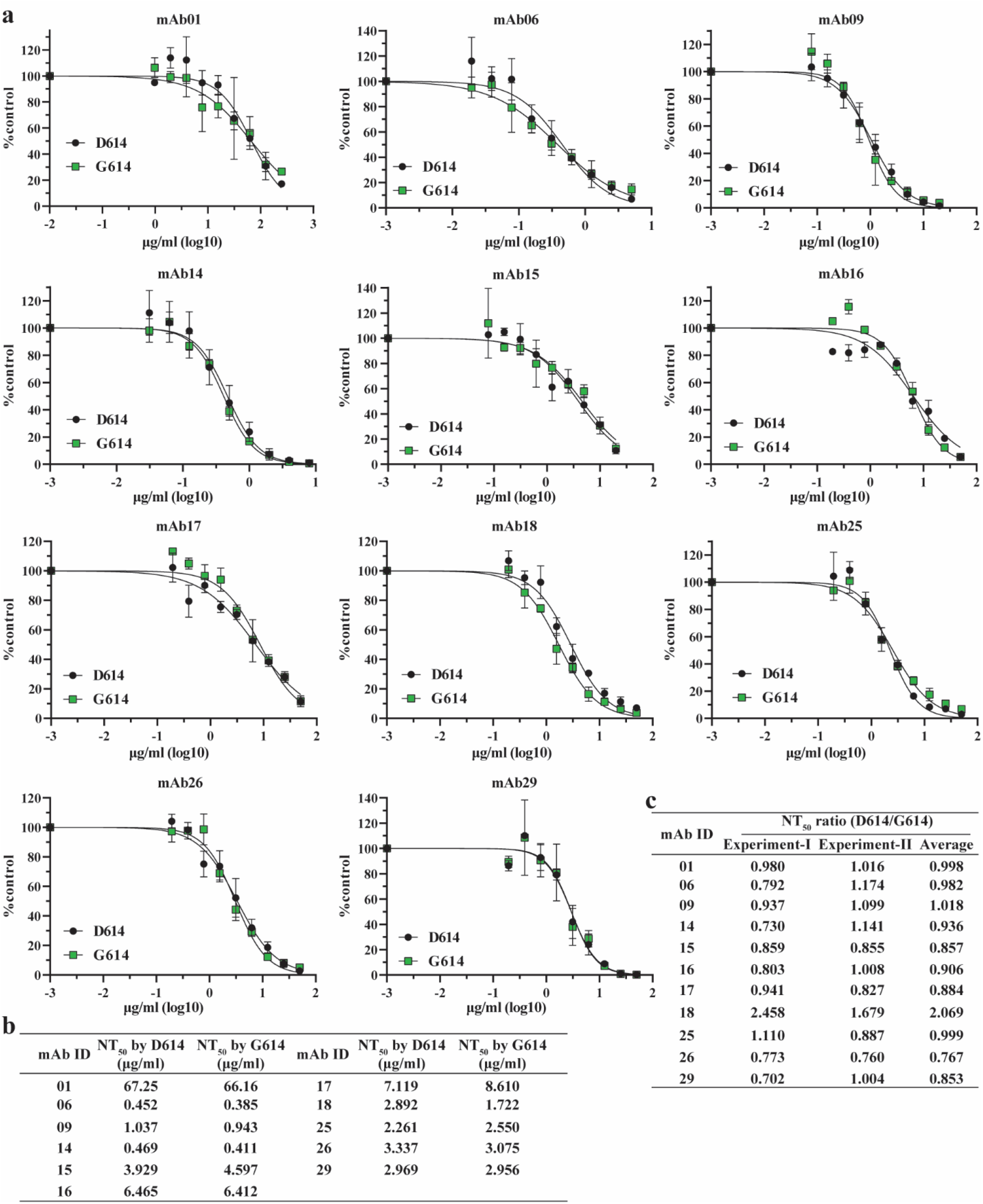
Neutralization activities of human mAbs against D614 and G614 mNeonGreen SARS-CoV-2 in Experiment II. **(a)** Neutralizing curves of eleven mAbs against D614 and G614 reporter SARS-CoV-2. Experiments were performed in replicates. The mean and standard deviations are shown. **(b)** Calculated NT_50_ values for all eleven mAbs. (c) Summary of NT_50_ ratios from two independent experiments. The ratios were determined by (D614 NT_50_)/(G614 NT_50_).

**Extended Data Table 1.**
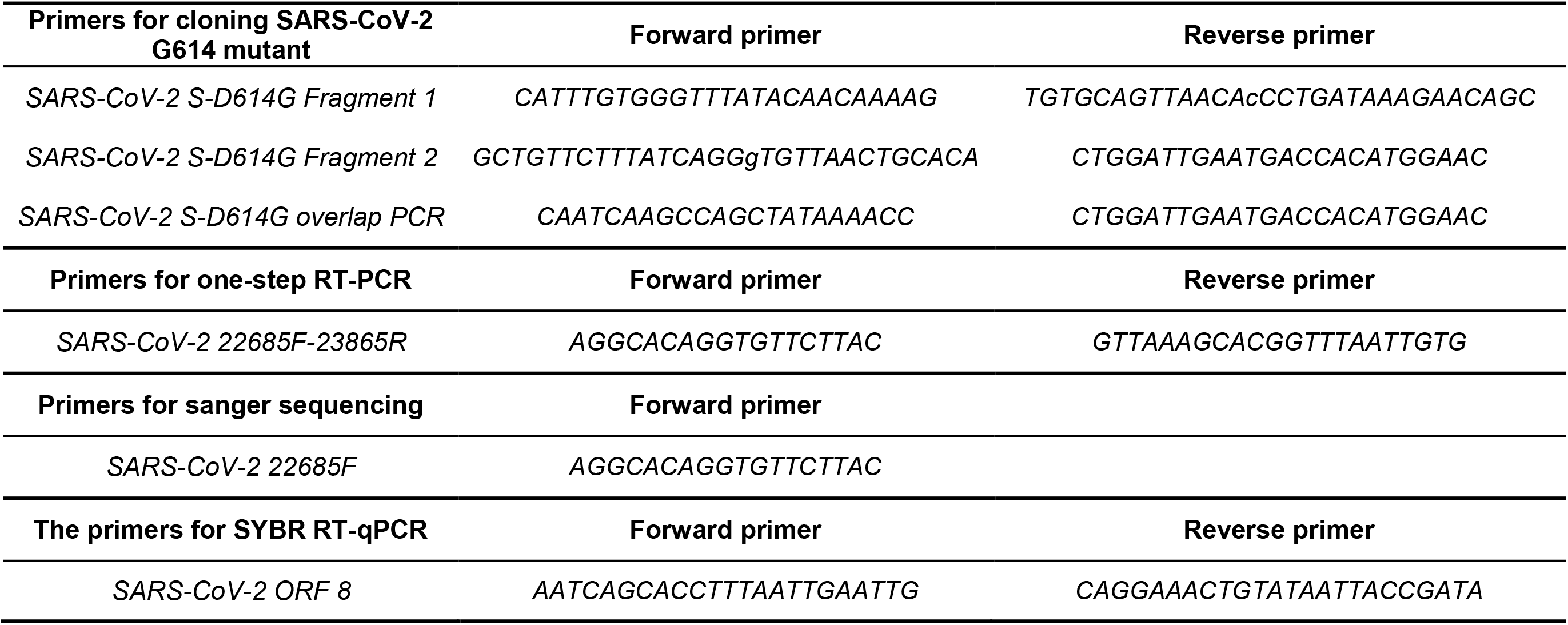
Primers for gene cloning and qPCR

